# Engineering a GelMA-dECM-Based 3D Bioprinted Liver Fibrosis Model: Methotrexate-Induced Functional and Molecular Validation

**DOI:** 10.1101/2025.08.09.669455

**Authors:** Mrunmayi Gadre, Kirthanashri S Vasanthan

## Abstract

Three-dimensional (3D) bioprinting represents a cutting-edge advancement in additive manufacturing, offering unprecedented precision in fabricating *in vitro* models that recapitulate native tissue architecture and function. In this study, we engineered a 3D bioprinted hepatic construct using a composite bioink comprising Gelatin Methacryloyl (GelMA), decellularized extracellular matrix (dECM) derived from rat liver, and HepG2 cells. GelMA was synthesized in-house to provide mechanical integrity and biocompatibility, while the liver-derived dECM offered essential biochemical cues to mimic the native hepatic microenvironment. The combination of GelMA and dECM uniquely provides mechanical robustness and bioactive cues essential for hepatic tissue mimicry. This synergy enhances cellular functionality and supports accurate fibrosis modeling for translational research. The bioprinted constructs were crosslinked using microbial transglutaminase and a photoinitiator to achieve structural stability.

Following fabrication, the 3D bioprinted hepatic constructs were evaluated using cytocompatibility assays (MTT, live/dead), functional assays (albumin, urea, LDH, ALT, and ALP secretion), and gene expression profiling to validate liver-specific function. Subsequently, liver fibrosis was induced in the 3D bioprinted constructs via methotrexate (MTX) exposure, and the fibrotic phenotype was confirmed through functional decline and upregulation of fibrosis-related genes. This study demonstrates a robust and physiologically relevant 3D bioprinted *in vitro* model of methotrexate-induced liver fibrosis, offering a valuable platform for translational applications in drug screening and hepatic disease modelling.

## 1.0 INTRODUCTION

Three-dimensional (3D) bioprinting represents a sophisticated advancement in additive manufacturing, enabling precise fabrication of 3D constructs derived from computer-aided designs (CAD) [1,2]. This technology necessitates the integration of viable cells and bioactive materials, known as bioinks, which facilitate the development of in vitro tissue models capable of mimicking the intricate physical and biochemical properties of native tissues. The core components of the bioprinting process include bioink formulation, selection of biomaterials, and advanced bioprinting techniques. The workflow involves three primary stages: pre- bioprinting, which includes creating a G-CODE based model and material selection; bioprinting, involving precise layer-by-layer deposition of bioinks; and post-bioprinting, focused on cellular maturation, proliferation, and functional validation of the fabricated constructs [3–6].

The biomaterials selected for bioink formulations are typically chosen based on their capacity to support key cellular processes such as adhesion, proliferation, differentiation, and migration [7–9]. Gelatin, a naturally derived polymer obtained through collagen denaturation, possesses intrinsic arginine-glycine-aspartic acid (RGD) motifs crucial for cellular attachment and communication. However, gelatin alone is biodegradable and mechanically unstable over extended periods. To overcome these limitations, GelMA is synthesized by chemically modifying gelatin with methacrylic anhydride, resulting in enhanced mechanical stability and improved longevity. GelMA scaffolds exhibit suitable mechanical properties, excellent biocompatibility, and non-toxicity, thus widely explored in diverse tissue engineering applications, including the culture of stem cells, hepatocytes, and cartilage regeneration [10].

dECM obtained from native liver tissues, provides essential biochemical cues to replicate native organ-specific microenvironments. The decellularization process aims to effectively eliminate cellular components while preserving critical ECM components such as collagens, elastin, fibrillin, fibronectin, laminin, proteoglycans (including heparan sulphate, chondroitin sulfate, keratan sulfate, and glycosaminoglycans (GAGs)), and embedded growth factors [11]. Various decellularization strategies including physical, chemical, and enzymatic are employed, with rigorous evaluations performed to ensure effective removal of cellular remnants, thereby guaranteeing the cytocompatibility and structural integrity of the resulting dECM [12–14]. The established criteria for successful decellularization include dsDNA content of less than 50 ng per mg ECM dry weight [15], DNA fragment lengths below 200 bp [16], and the absence of visible nuclear materials verified through DAPI or Haematoxylin and Eosin (H&E) staining [17]. In this study, chemical decellularization via EDTA and sequential concentrations of sodium dodecyl sulfate (SDS) is utilized to produce liver-derived dECM.

Current *in vitro* liver fibrosis models often lack physiological complexity, functional robustness, and fail to adequately recapitulate the dynamic molecular events observed in vivo, limiting their translational applicability [18–20]. The present study addresses these gaps by combining GelMA, providing mechanical stability [21], and liver-derived dECM, supplying native biochemical signalling cues [22, 23], to engineer a bioink specifically tailored for hepatic tissue engineering. Incorporation of HepG2 cells within this bioink facilitates the generation of liver constructs with improved structural and functional fidelity, enabling precise modeling of non-fibrotic and fibrotic liver conditions [24–26]. This innovative model not only enhances the physiological relevance and predictive accuracy for studying hepatic fibrosis but also significantly advances translational applications, including drug screening, toxicity evaluations, and disease modeling, ultimately accelerating the discovery of effective therapeutic interventions [27, 28].

## 2.0 MATERIALS AND METHODS

### 2.1 Materials

Dulbecco’s modified Eagle’s medium (DMEM, high glucose w/ 4.5gms Glucose per litre, L-Glutamine and Sodium bicarbonate w/o Sodium pyruvate and Dulbecco’s phosphate-buffered saline (DPBS) was procured from Himedia, India. Fetal bovine serum (FBS) was got from Gibco. Penicillin/streptomycin, trypsin, trizol and methotrexate were ordered from Sigma Aldrich and Albumin (BCG) [ab235628] Assay Kit from Abcam. Lactate Dehydrogenase (LDH) activity [E-BC-K046-M] assay kit and Urea (BUN) Calorimetric based assay (Urease method) [E-BC-K183-M] were purchased from Elab sciences. The Alanine Transaminase (ALT) [ab241035] and alkaline phosphatase (ALP) [ab83369] calorimetric kit-based assay from Abcam. Sodium dodecyl sulfate, phenol, chloroform, isoamyl alcohol, isopropanol and papain were purchased from Sisco Research Laboratories, India. Ethylenediamine tetra acetic acid and formaldehyde was bought from Qualigens, India. Glutaraldehyde was ordered from Sigma Aldrich, India. Proteinase K was purchased from Invitrogen, India. Coomassie brilliant blue and protease inhibitor cocktail was obtained from Merck, India. Sodium acetate was purchased from Tokyo Chemical Industry, India. RIPA buffer was procured from Himedia, India. Pierce™ CA Protein Assay Kit was purchased from Thermo Fisher Scientific, India and Blyscan™ - sulfated Glycosaminoglycan (sGAG) assay kit from biocolor, United Kingdom, PrimeScript™ 1st strand cDNA Synthesis Kit and TB Green Premix Ex Taq (Tli RNase H Plus) was brought from Takara, USA.

### 2.2 Animal ethics approval

All the animals used in this study were approved by the Institutional Animal Ethics Committee (IAEC), Manipal Academy of Higher Education, Manipal which is in accordance with CPSCEA guidelines (IAEC/KMC/121/2022). All the experiments involving the animals were performed following ARRIVE guidelines.

### 2.3 METHODS

#### 2.3.1 Cell Culture

Human hepatocellular carcinoma (HepG2) cells were procured from the National Centre for Cell Science (NCCS), Pune, India. Cells were cultured in Dulbecco’s Modified Eagle Medium (DMEM) supplemented with 10% (v/v) fetal bovine serum (FBS) and 1% (v/v) penicillin-streptomycin (P/S). Cultures were maintained at 37°C in a humidified atmosphere with 5% CO₂. Cells were routinely sub-cultured upon reaching 80–90% confluence using standard trypsinization protocols.

#### 2.3.2 Liver Harvest and Decellularization Protocol

Male Wistar rats (150–250 g; 9–12 weeks old) were anesthetized via intraperitoneal injection of sodium pentobarbital (50 mg/kg). The excised liver was rinsed with sterile distilled water and immersed in 0.001 M ethylenediaminetetraacetic acid (EDTA) for 30 minutes to facilitate initial matrix loosening. Post-treatment, the liver was sectioned into small fragments and perfused with 60 mL of distilled water to remove blood residues.

Decellularization was carried out following an optimized protocol previously reported by authors [29]. Briefly decellularization was carried through sequential perfusion with sodium dodecyl sulfate (SDS) at increasing concentrations (0.1%, 0.25%, 0.5%, 0.75%, and 1%, 60 mL each per concentration), introduced through 8–10 injection sites per tissue fragment to ensure uniform exposure [29]. Upon completion, the tissue was rinsed extensively with distilled water until a translucent appearance was observed, indicating successful decellularization. The resulting decellularized extracellular matrix (dECM) was subjected to morphological and biochemical characterization, including DNA quantification, histological staining, protein profiling, and ultrastructural imaging via scanning electron microscopy (SEM).

#### 2.3.3 Histological Evaluation

Native and decellularized liver tissues were fixed in 10% neutral-buffered formalin and stored at ambient temperature until further processed by microtome method [30]. The 3D bioprinted scaffold were covered with optimal cutting temperature cryogel (Sakura Finetek, USA) and cryosectioning was performed in Cryostat RWD Minux FS800 (Crystal bio equipment). All the sections were stained in house, briefly the sections were washed with distilled water for 60 seconds, and the slides were dipped in haematoxylin for 10 minutes and washed under tap water for 5 minutes. Further the slides were subjected to eosin for 20 seconds and washed with 75% ethanol, followed by 90% and 100%. Once the slides were dried completely, they were fixed with Dibutylphthalate Polystyrene Xylene (DPX). Stained sections were visualized using an Olympus CX43 microscope (Olympus, Japan), and images were captured for histopathological comparison.

#### 2.3.4 Scanning Electron Microscopy

To assess ultrastructural preservation, tissue samples were fixed in 2.5% glutaraldehyde at 4°C for 2 hours. The fixed samples were dehydrated through a graded ethanol series (10%–100%) with 10-minute intervals per step, followed by overnight drying at 37°C [31]. SEM imaging was conducted using the ZEISS EVO MA18 scanning electron microscope (Carl Zeiss, Germany) at the Central Instrumentation Facility, Manipal Institute of Technology, MAHE.

#### 2.3.5 Genomic DNA Isolation and Quantification

Genomic DNA was isolated from 50 mg of native and decellularized liver tissues using a phenol:chloroform:isoamyl alcohol extraction method (n = 3). Tissue homogenization was performed in 1× PBS using a Scilogex SCI16 PRO homogenizer (USA), followed by centrifugation at 5000 rpm for 10 minutes. The supernatant was discarded, and the pellet was treated with 200 µL of lysis buffer, 400 µL of phenol, and 10 µL of Proteinase K. After mixing via rotospin (25 rpm, 10 minutes), samples were incubated at 55°C for 2 hours in a water bath.

Subsequent phase separation was performed using sequential extractions with phenol and a 1:1 mixture of chloroform:isoamyl alcohol (CIA), followed by centrifugation at 8000 rpm for 10 minutes after each extraction. The aqueous layer was collected and treated with chilled isopropanol and sodium acetate to precipitate DNA. The pellet was washed twice with 100% ethanol and dissolved in 1× TE buffer after air drying. The purity and yield of DNA were confirmed via 1% agarose gel electrophoresis and spectrophotometric analysis.

#### 2.3.6 Quantification of Total Protein and Glycosaminoglycans

Total protein content was quantified using the Bicinchoninic Acid (BCA) assay. Proteins were extracted from 50 mg of both native and decellularized tissues using 1× radioimmunoprecipitation assay (RIPA) buffer supplemented with protease inhibitor cocktail. Equal volumes of the lysate and working reagent from the BCA kit were mixed and incubated at 37°C for 30 minutes. Absorbance was measured at 562 nm, and concentrations were interpolated from a standard curve.

The extracted protein samples were further resolved by sodium dodecyl sulfate– polyacrylamide gel electrophoresis (SDS-PAGE) using a 6% resolving and 4% stacking gel under constant voltage (100 V for 90 minutes). Gels were stained with Coomassie Brilliant Blue and imaged using the Invitrogen iBright CL1500 imaging system (Thermo Fisher Scientific).

#### 2.3.7 Gelatin Methacryloyl Synthesis

GelMA was synthesized following an optimized protocol previously reported [32]. Briefly, gelatin type A (porcine, Bloom strength 175 g) was dissolved at a concentration of 10% (w/v) in phosphate-buffered saline (PBS, pH 7.4) at 50°C. Methacrylic anhydride (MA) was added (0.6 g MA per 1 g gelatin) under vigorous stirring (1200 rpm) at 50°C for 1 hour. The resultant mixture was centrifuged at 4000 rpm for 2 minutes to remove excess MA. The pellet was discarded, and the supernatant was diluted with an equal volume of PBS, followed by dialysis against deionized water using a dialysis membrane (10 kDa MWCO, Hi-Media, India) at 40°C for 5 days, changing the water daily. The purified GelMA solution was lyophilized and stored at -20°C until use. GelMA was characterized in comparison to native gelatin using Fourier-transform infrared spectroscopy (FTIR), which confirmed methacrylation through the detection of characteristic amide bond modifications.

#### 2.3.8 Decellularized ECM Solubilization

Decellularized liver tissue comprises extracellular matrix proteins such as collagen, elastin, fibrillin, fibronectin, laminin, proteoglycans (heparan sulfate, chondroitin sulfate, keratan sulfate, glycosaminoglycans), and embedded growth factors. Prior to incorporation into bioink formulations, dECM was solubilized by lyophilization for 72 hours, followed by milling into fine powder. The powder (2 mg/mL) was enzymatically digested with pepsin (10% w/w) in 0.5 N acetic acid at room temperature under constant stirring for 48 hours. Post-digestion, the solution was centrifuged (3000 rpm, 15 min), neutralized with 10 N sodium hydroxide, and quantified using BCA assay (Invitrogen). Protein integrity in solubilized dECM compared to native tissue was assessed by SDS-PAGE.

#### 2.3.9 Computer-Aided Design (CAD) for Bioprinting

CAD models for bioprinting were generated using Openscad software and converted into stereolithography (STL) files. Model specifications, including dimensions and extrusion parameters, were defined precisely within the software. Bioprinting requires digital models typically obtained from imaging modalities (X-ray, computed tomography [CT], magnetic resonance imaging [MRI]) or designed directly through CAD software. All generated STL files underwent validation using computer-aided manufacturing (CAM) software prior to printing, ensuring compatibility and printability [33–36]. Bioink extrusion volumes were accurately calculated based on layer dimensions specified in STL files.

#### 2.3.10 Cytotoxicity Analysis of Solubilized dECM and GelMA on HepG2 Cells

To evaluate potential cytotoxic effects of bioink components, solubilized dECM was added to HepG2 cell cultures at varying densities (10 µL dECM per 100 µL culture medium). Cell viability was assessed at 72 hours using the MTT assay. After incubation, media was removed, cells were briefly washed with PBS, and 100 µL of 0.5% MTT solution was added per well, incubated for 2 hours in darkness at 37°C, followed by removal of MTT solution. Formazan crystals were dissolved by adding 100 µL DMSO per well, and absorbance readings were recorded at 570, 590, and 630 nm.

#### 2.3.11 Characterization of 3D Bioprinted Constructs

GelMA-based bioink was formulated by combining lyophilized GelMA with Lithium Phenyl- 2,4,6-trimethyl-benzoylphosphinate (LAP; Sigma Aldrich, China), PBS, microbial transglutaminase (MTgase), and HepG2 cells. Bioink was loaded into sterile syringes and extruded through a dual-extrusion bioprinter (Alfatek Systems, India) fitted with a 0.7 mm diameter needle at pressures of 120–180 kPa and deposition speed of 600 mm/min at room temperature. Constructs (1 × 1 cm) were printed in a rectilinear grid pattern and crosslinked using UV illumination in the presence of LAP.

##### 2.3.11.1 Degradation Study

Scaffold degradation was monitored by measuring initial wet weights, intermediate wet weights, and post-lyophilization dry weights at defined intervals. Structural changes and patterns on constructs were recorded using microscopy throughout the incubation period.

##### 2.3.11.2 Induction of Fibrosis

Fibrotic conditions were induced in HepG2 cells cultured on bioprinted constructs by exposure to 10 mM methotrexate, optimized from previous studies, over a 72-hour period. Post-treatment, cell viability was analyzed by MTT assay, and morphological evidence of fibrosis was evaluated using H&E staining. Cells were fixed in formaldehyde, sequentially dehydrated using graded ethanol washes, stained, and mounted with DPX prior to microscopic examination.

##### 2.3.11.3 Live/Dead Viability Assay

To corroborate MTT findings, viability of HepG2 cells post-fibrosis induction was assessed via live/dead fluorescent staining. After removing culture media and washing with Dulbecco’s phosphate-buffered saline (DPBS), cells were incubated with a fresh live/dead reagent mixture (100 µL for 96-well plates or 200 µL for 24-well plates) in the dark at room temperature for 45 minutes. Fluorescent imaging was performed, and three-dimensional cell distributions were acquired using confocal Z-stack microscopy.

#### 2.3.12 Biochemical Assays

The biochemical markers including albumin, urea, lactate dehydrogenase (LDH), alkaline phosphatase (ALP), and alanine aminotransferase (ALT) were quantified from conditioned culture media using respective colorimetric assay kits, following the manufacturers’ protocols. These markers were chosen based on their relevance to liver-specific functionality.

##### 2.3.12.1 Albumin Assay

Albumin levels were determined using a colorimetric assay kit based on the interaction of Bromocresol Green (BCG) with albumin, forming a chromophore measurable at 620 nm. In brief, 50 µL of culture supernatant was mixed with 100 µL of the reagent mix and incubated at room temperature for 25 minutes. The absorbance was measured at 620 nm using a microplate reader, and albumin concentration was quantified using a standard calibration curve.

##### 2.3.12.2 Urea Assay

Urea concentration was measured using a urease-based colorimetric kit, which catalyses the hydrolysis of urea to ammonia and carbon dioxide. The released ammonia reacts with a chromogenic reagent to produce a green-coloured complex detectable at 580 nm. A 4 µL sample aliquot was mixed with 50 µL of enzyme reagent and incubated at 37°C for 10 minutes.

Subsequently, 125 µL each of Reagents 4 and 5 were added, followed by a second 10-minute incubation. Absorbance was measured at 580 nm, and urea concentrations were calculated from a standard curve.

##### 2.3.12.3 Alkaline Phosphatase (ALP) Assay

ALP activity was evaluated using a high-throughput-compatible kit (Abcam) based on the hydrolysis of p-nitrophenyl phosphate (pNPP) to p-nitrophenol (pNP), detectable at 405 nm. Briefly, 50 µL of 5 mM pNPP was added to each well containing sample, followed by 10 µL of ALP enzyme solution in the standard wells. The reaction was incubated at 25°C for 60 minutes, stopped using 20 µL of stop solution, and the absorbance was measured at 405 nm. The ALP concentration was extrapolated from a standard curve.

##### 2.3.12.4 Alanine Aminotransferase Assay

ALT activity was assessed using a colorimetric ALT activity assay kit (Abcam). In brief, 0.5–2.5 µL of sample was adjusted to 5 µL with assay buffer. A freshly prepared 25 µL reaction mix was added to each well containing samples, standards, and controls. An additional 25 µL of background mix was added to background control wells. After incubation, the absorbance was read at 570 nm, and the ALT activity was calculated using a standard curve.

##### 2.3.12.5 Lactate Dehydrogenase Assay

LDH activity was measured using a colorimetric kit based on the enzymatic conversion of lactic acid to pyruvic acid in the presence of coenzyme I. The resulting pyruvate forms a brownish-red hydrazone derivative with dinitrophenylhydrazine under alkaline conditions, detectable at 450 nm. Briefly, 200 µL of sample was mixed with 250 µL of substrate buffer and 50 µL of coenzyme I, followed by incubation at 37°C for 15 minutes. Then, 250 µL of chromogenic agent was added and incubated again for 10 minutes. Finally, 2500 µL of alkaline developer was added, and absorbance was recorded at 450 nm. LDH concentration was calculated using a standard curve.

#### 2.3.13 Gene Expression Analysis

##### 2.3.13.1 RNA Isolation

Scaffold samples were enzymatically digested using 1 mL of collagenase type I at 37°C for 15–20 minutes. The digested material was centrifuged at 10,000 rpm for 5 minutes at 4°C, washed with 1× PBS, and centrifuged again. The pellet was treated with 800 µL of TRIzol reagent followed by 160 µL of chloroform, vortexed, and incubated at room temperature for 2 minutes. After centrifugation at 12,000 rpm for 20 minutes at 4°C, the upper aqueous phase was transferred to fresh tubes, and an equal volume of molecular-grade isopropanol was added. Following incubation and centrifugation at 13,000 rpm for 20 minutes, the RNA pellet was washed twice with ice-cold 70% ethanol and centrifuged at 7500 rpm for 5 minutes. The pellet was air-dried and resuspended in 10 µL of DEPC-treated water. RNA quantity was assessed using a microplate reader, and samples were stored at –80°C.

##### 2.3.13.2 cDNA Synthesis

cDNA synthesis was performed using a commercial reverse transcription kit. The master mix was prepared as per the manufacturer’s protocol, and 7 µL of the mix was aliquoted into sterile tubes along with the required volumes of RNA and dilution buffer. The thermal cycling conditions were 37°C for 30 minutes, 85°C for 5 seconds, and hold at 4°C. Synthesized cDNA was stored at –20°C.

##### 2.3.13.3 Quantitative Real-Time PCR

qRT-PCR was carried out using a commercial SYBR Green-based kit. Primer and template master mixes were prepared according to kit instructions. Each reaction contained 6 µL of primer mix and 4 µL of template mix. For the non-template control (NTC), 4 µL of DEPC water was added instead of the template. The thermal cycling conditions were:

- 95°C for 30 seconds (initial denaturation)
- 40 cycles of 95°C for 5 seconds and 60°C for 34 seconds (amplification)
- Melting curve analysis: 95°C for 15 seconds, 60°C for 1 minute, and 95°C for 15 seconds.

Post-run, reactions were stored at –20°C for downstream analysis. Primer sequences are provided in the **Table 1**.

**Table 1.**
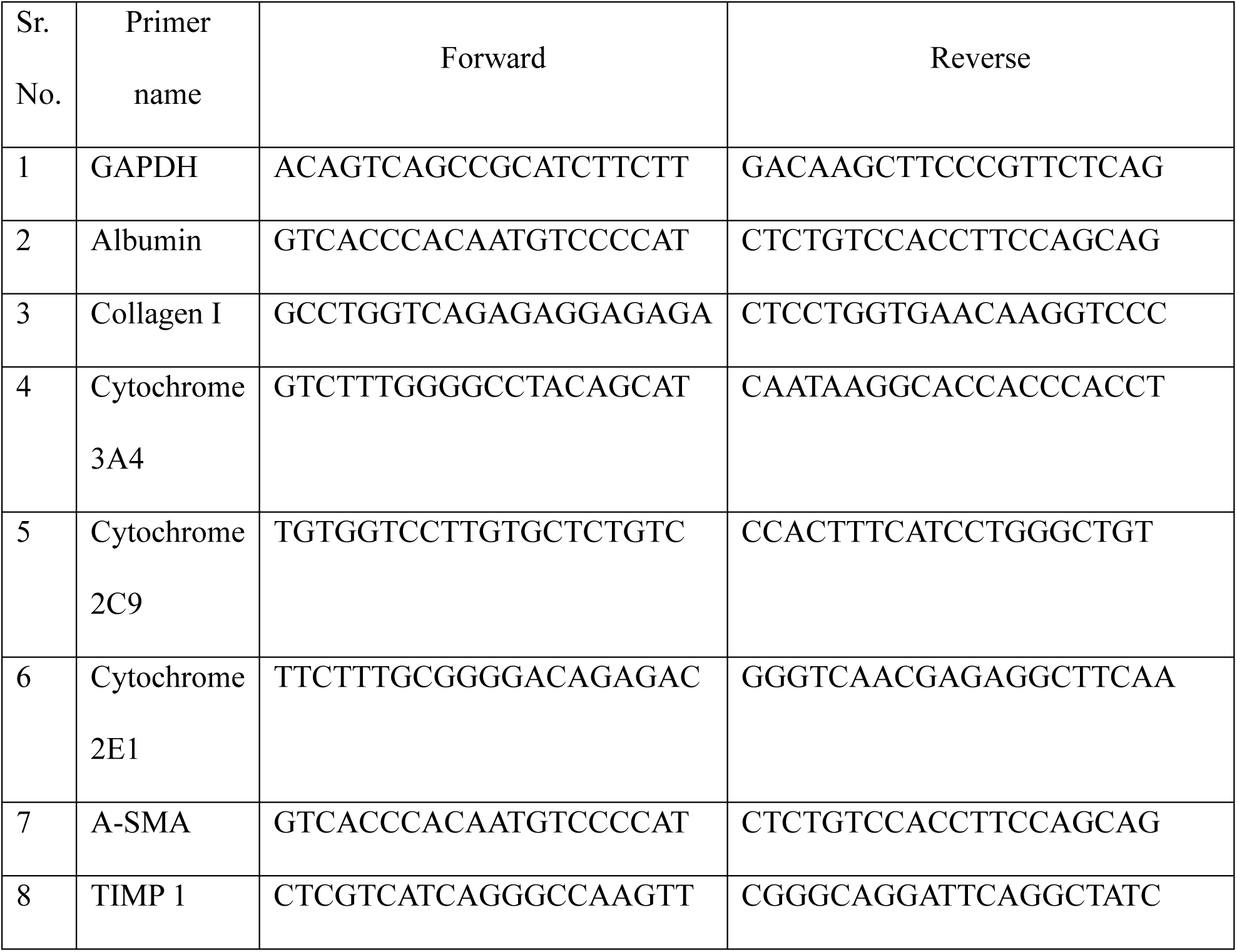
The forward and reverse primer sequences.

### 3.0 RESULTS AND DISCUSSION

#### 3.3.1 Liver Harvesting and Decellularization Outcomes

The decellularization of rat liver tissue was successfully accomplished within approximately 12 hours utilizing a stepwise chemical approach. Three freshly harvested rat livers were dissected into uniformly and sequentially exposed to EDTA, followed by graded concentrations of SDS (0.1%, 0.25%, 0.5%, 0.75%, and 1%). This controlled treatment effectively facilitated the removal of cellular components, as confirmed by visual assessment (**Figure 1**). The native liver tissue initially exhibited a dark red appearance (**Figure 1A)**, progressively transitioning to a translucent state indicative of successful cell removal upon completion of the SDS treatment series (**Figure 1G**).

**Figure 1.**
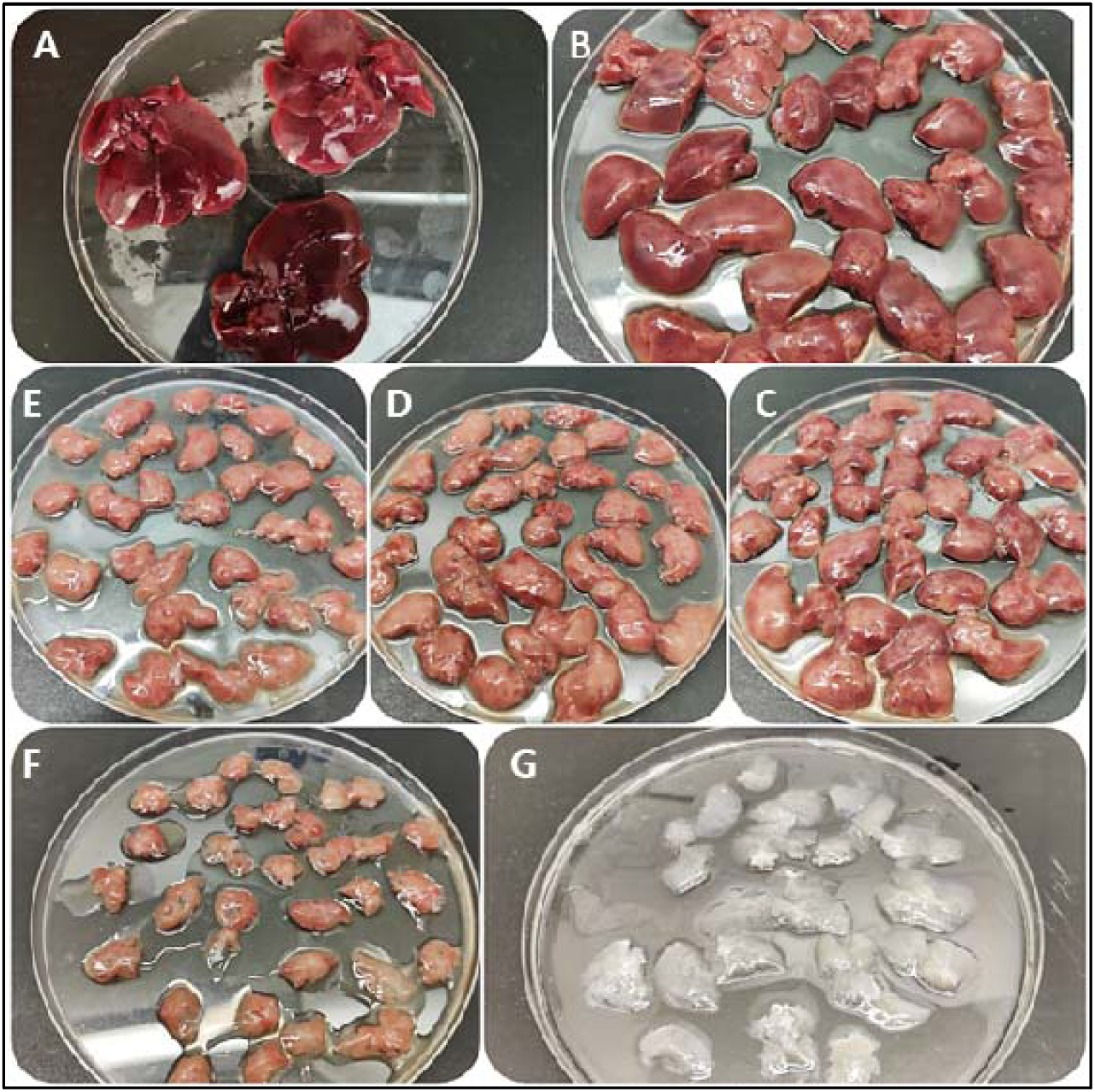
Rat Liver Decellularization A – native liver; B – EDTA; C to G – gradually increasing concentrations of SDS (0.1%, 0.25%, 0.5%, 0.75% and 1%)

Post-decellularization, liver matrices were stored at -80 °C in PBS containing sodium azide until subsequent analyses. Comparative analysis revealed a significant reduction in liver mass from pre-treatment to post-decellularization, resulting in dECM weight of approximately 1.1 g. Quantitatively, the average mass reduction from native liver tissue was approximately 87 ± 0.003%. Although weight reduction alone is insufficient to conclusively demonstrate complete decellularization, the consistent and substantial reduction observed aligns with effective cellular removal (**Table 2**). These findings were further substantiated through comprehensive biochemical and ultrastructural characterization studies.

**Table 2.**
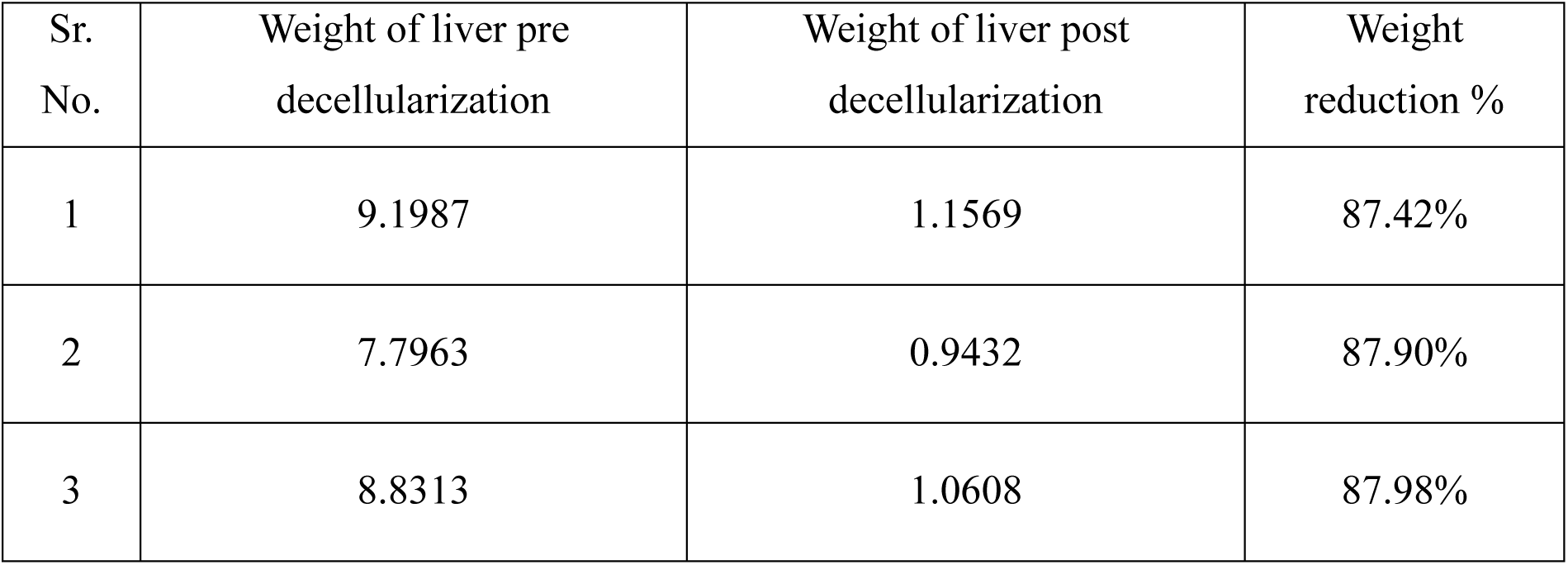
Rat Liver weight pre and post decellularization.

#### 3.3.2 Histological Analysis

H&E staining was performed to evaluate the efficacy of decellularization and the preservation of the ECM structure. Microscopic examination revealed complete removal of cellular components from the dECM, as demonstrated by the absence of haematoxylin-positive nuclear staining (**Figure 2B**). The dECM predominantly displayed eosin-positive staining (pink), indicative of retained ECM components such as collagen fibres. In contrast, the native liver tissue exhibited intense haematoxylin staining (purple), confirming the presence of intact nuclei and cellular structures along with ECM components (**Figure 2A**). These histological findings substantiate successful decellularization, while maintaining the integrity of the liver ECM architecture.

**Figure 2.**
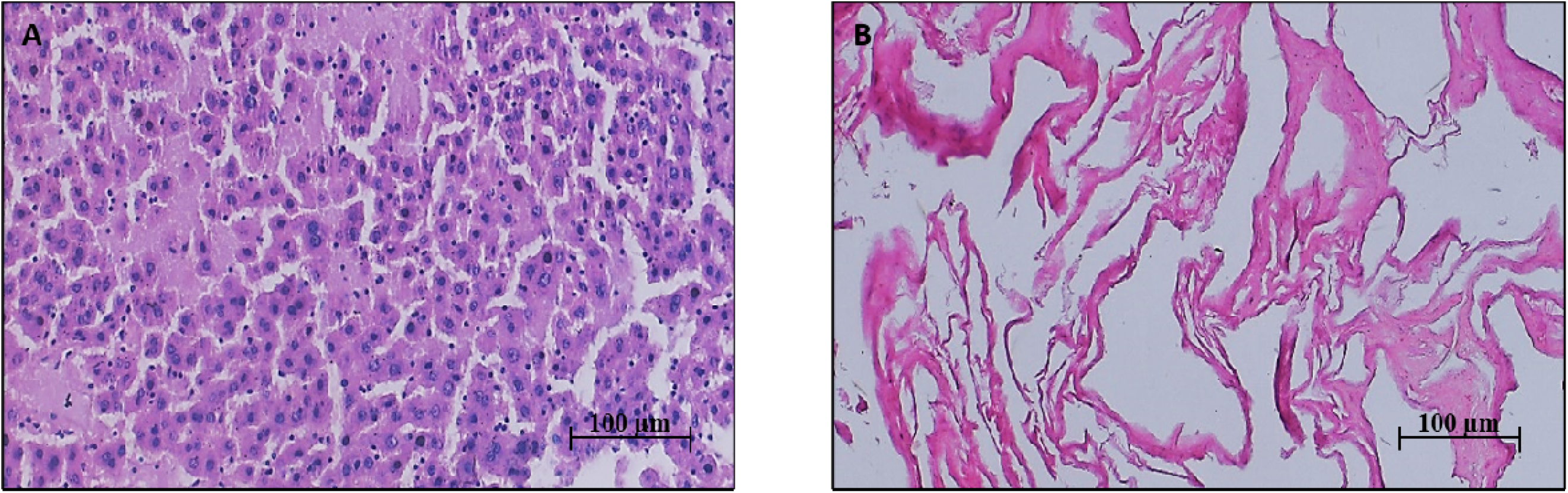
H&E staining of (A) native liver tissue; (B) decellularized liver tissue (40X magnification)

#### 3.3.3 Scanning Electron Microscopy Analysis

SEM analysis was conducted to examine the ultrastructural alterations in liver tissue before and after decellularization. Representative SEM micrographs illustrated substantial differences between native liver tissue and dECM, as depicted in **Figure 3**. Native liver tissue (**Figure 3A**) displayed a densely packed architecture characterized by smooth, spherical cellular structures, indicative of intact hepatocytes. Conversely, the decellularized liver scaffold (**Figure 3B**) exhibited a highly porous, fibrous ECM network devoid of cellular morphology, demonstrating successful cellular removal. These SEM findings confirm effective decellularization while preserving the essential ultrastructural integrity of the ECM, supporting its potential application as a scaffold for tissue engineering purposes.

**Figure 3.**
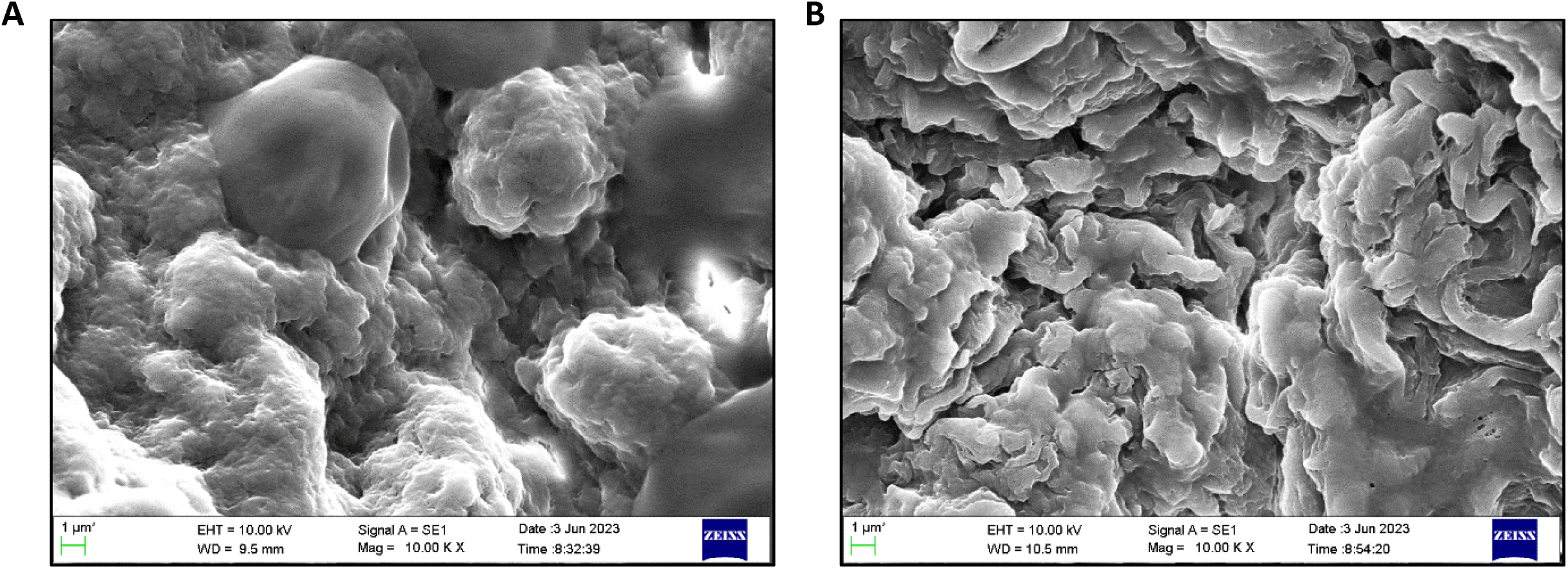
SEM images of (A) native liver tissue; (B) decellularized liver tissue

#### 3.3.4 DNA Quantification and Agarose Gel Electrophoresis

Quantitative and qualitative DNA assessments were conducted to confirm the effective removal of cellular components following decellularization. DNA content, normalized against the initial wet weight of each sample, demonstrated a significant reduction in the decellularized dECM compared to native liver tissue (**Figure 4A**). Specifically, the DNA content in native liver was quantified as 1200 ± 8.05 ng/mg, while the dECM exhibited substantially reduced residual DNA at 10 ± 2.54 ng/mg (values expressed as mean ± SD, n = 4 per group).

**Figure 4.**
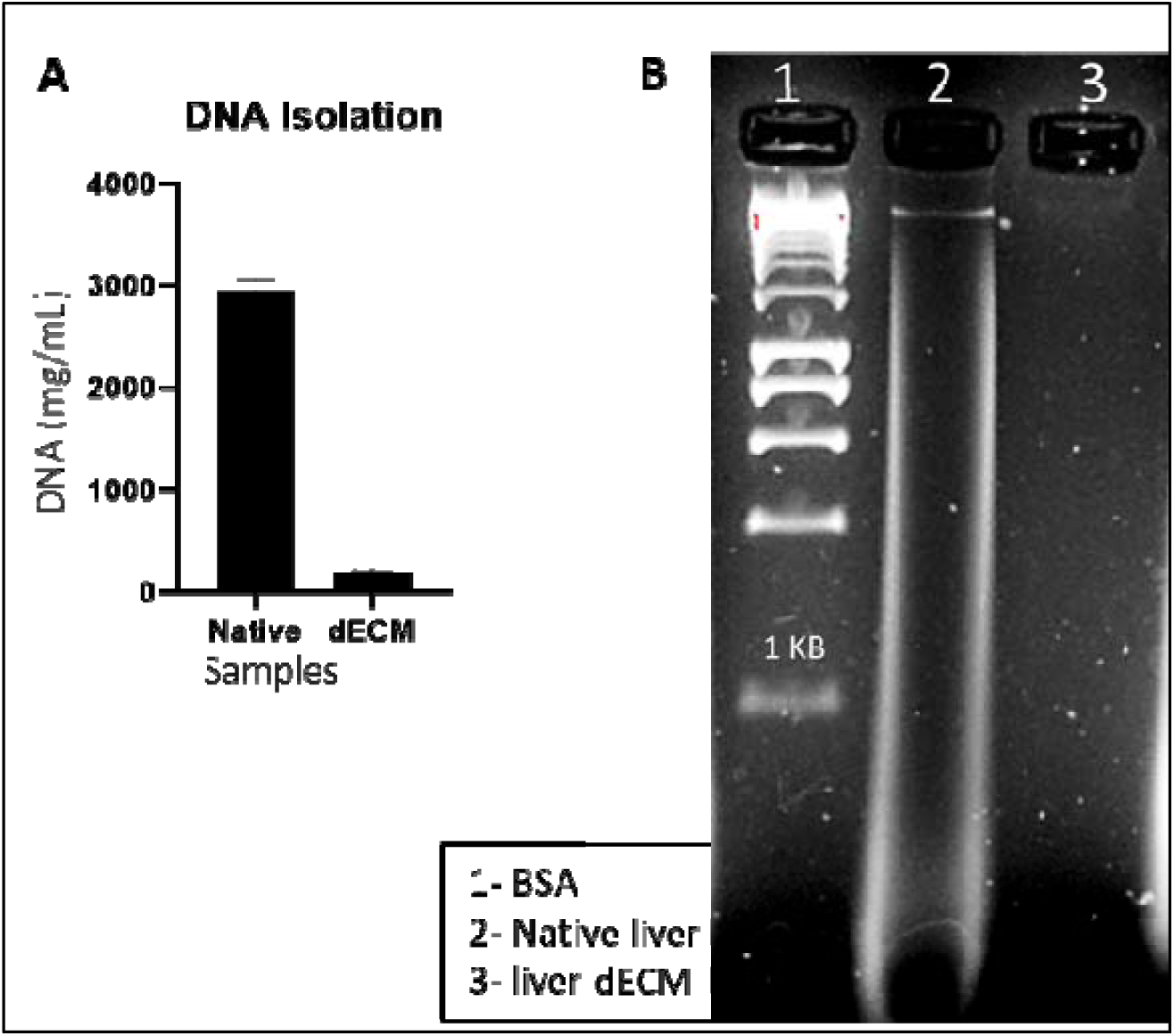
(A) DNA concentration post isolation from native and dECM; (B) 1% agarose gel; * : statistically significant (p<0.05) (B) 1% agarose gel

Further confirmation through agarose gel electrophoresis (**Figure 4B**) revealed prominent DNA bands in native liver samples, indicative of intact genomic material. In contrast, the absence of visible DNA bands in the dECM samples confirmed efficient genomic material removal, aligning with established standards for decellularization efficacy.

#### 3.3.5 Total Protein Quantification and SDS-PAGE Analysis

Protein integrity and composition in native liver, dECM, Gelatin, and GelMA were evaluated through SDS-PAGE analysis and BCA quantification assay (**Figure 5**). SDS-PAGE revealed distinct bands indicating the preservation of key extracellular matrix proteins, including the characteristic α1 and α2 chains of type I collagen at approximately 120 kDa and the β-chain at approximately 250 kDa (**Figure 5B**). Comparative analysis confirmed similar protein profiles between native liver and dECM samples, demonstrating effective retention of ECM proteins post-decellularization. Quantitatively, total protein concentrations measured via BCA assay were as follows: Gelatin, 1.20 ± 0.98 µg/mL; GelMA, 1.57 ± 1.20 µg/mL; native liver, 1.45 ± 0.67 µg/mL; and dECM, 1.28 ± 0.83 µg/Ml (**Figure 5A**). Results are presented as mean ± SD (n = 4 per group), suggesting minimal protein loss during the decellularization process.

**Figure 5.**
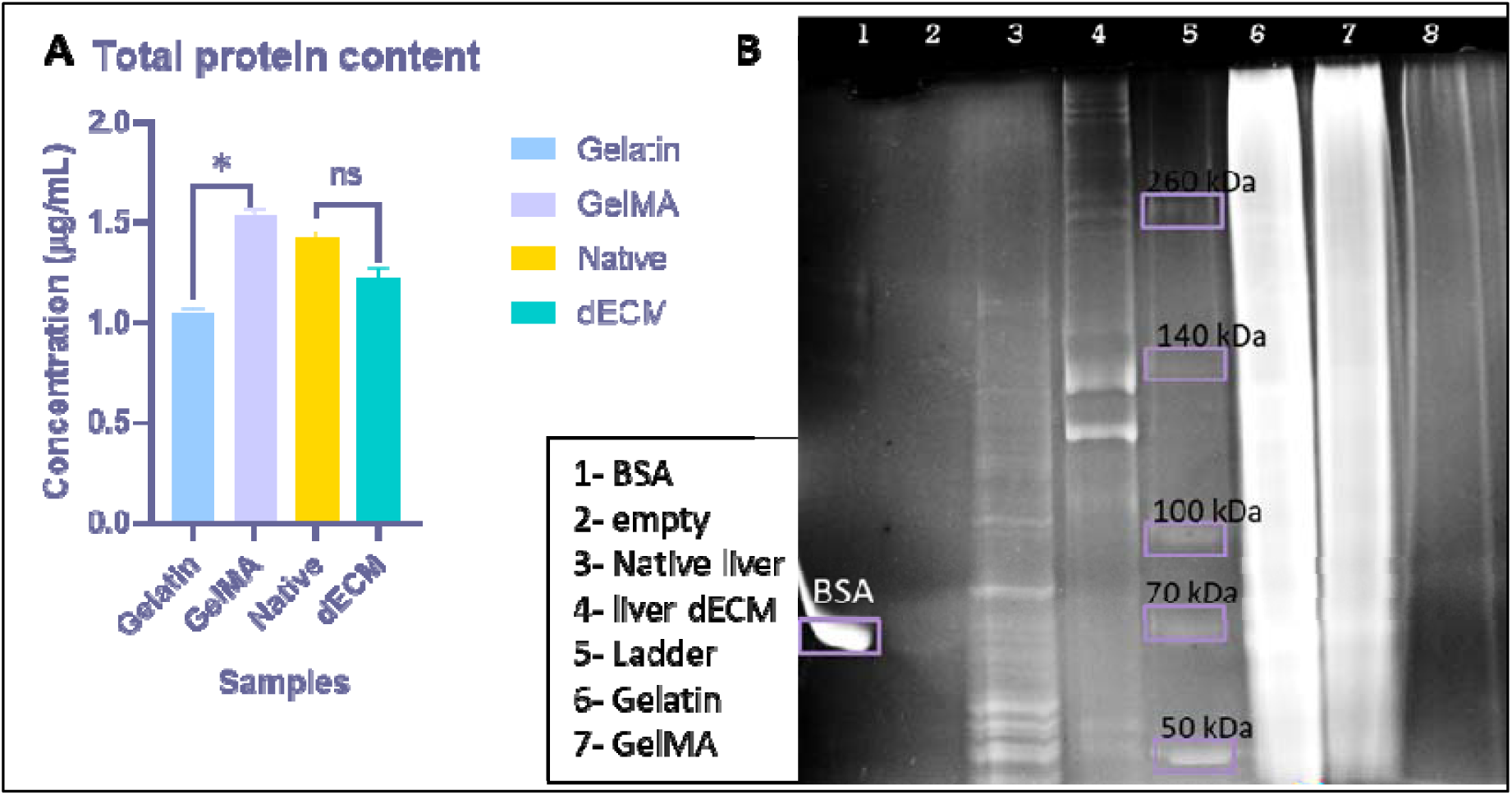
Total Protein concentration post isolation from native and dECM (A) Protein concentration * : statistically significant (p<0.05) and ns : non-significant; (B) SDS PAGE of isolated proteins

#### 3.3.6 Quantification of Glycosaminoglycans (GAGs)

The sGAG content in native and decellularized rat liver tissues was quantified to assess ECM preservation post-decellularization. **Figure 6** demonstrates a moderate retention of sGAGs in dECM compared to native tissue, with concentrations quantified at 12.56 ± 1.20 µg/mL and 14.98 ± 0.95 µg/mL, respectively (mean ± SD; n = 4 per group). This significant retention of sGAG indicates that the decellularization protocol preserved important ECM components, thereby enhancing the potential biological functionality of the dECM scaffold.

**Figure 6.**
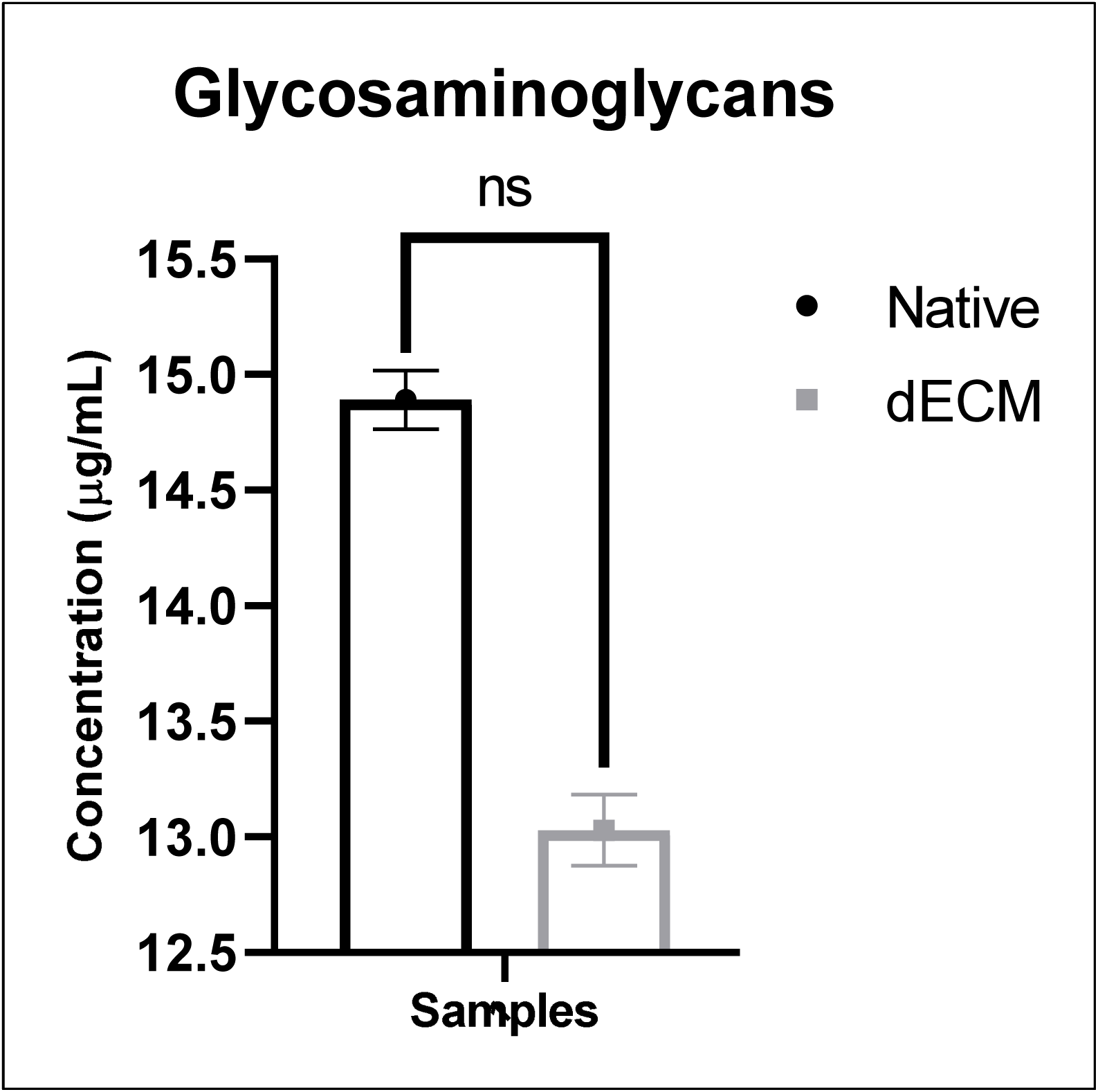
sGAG concentration post isolation from native and dECM; (p<0.05) ns : non-significant

#### 3.3.7 Gelatin Methacryloyl Synthesis

GelMA was synthesized and the characterization was performed using the FTIR and the presence of amide bonds were at different peaks. The peak at 1650 cm-1, 1550 cm-1 and 1450 cm-1 confirmed the presence of amide bond I, II and III respectively (**Figure 7**). The NMR was also performed and the degree of substitution was calculate to be 70% which was found sufficient for stability of GelMA. The other characterization of GelMA has already been studied and established in the patent filled by the research team.

**Figure 7.**
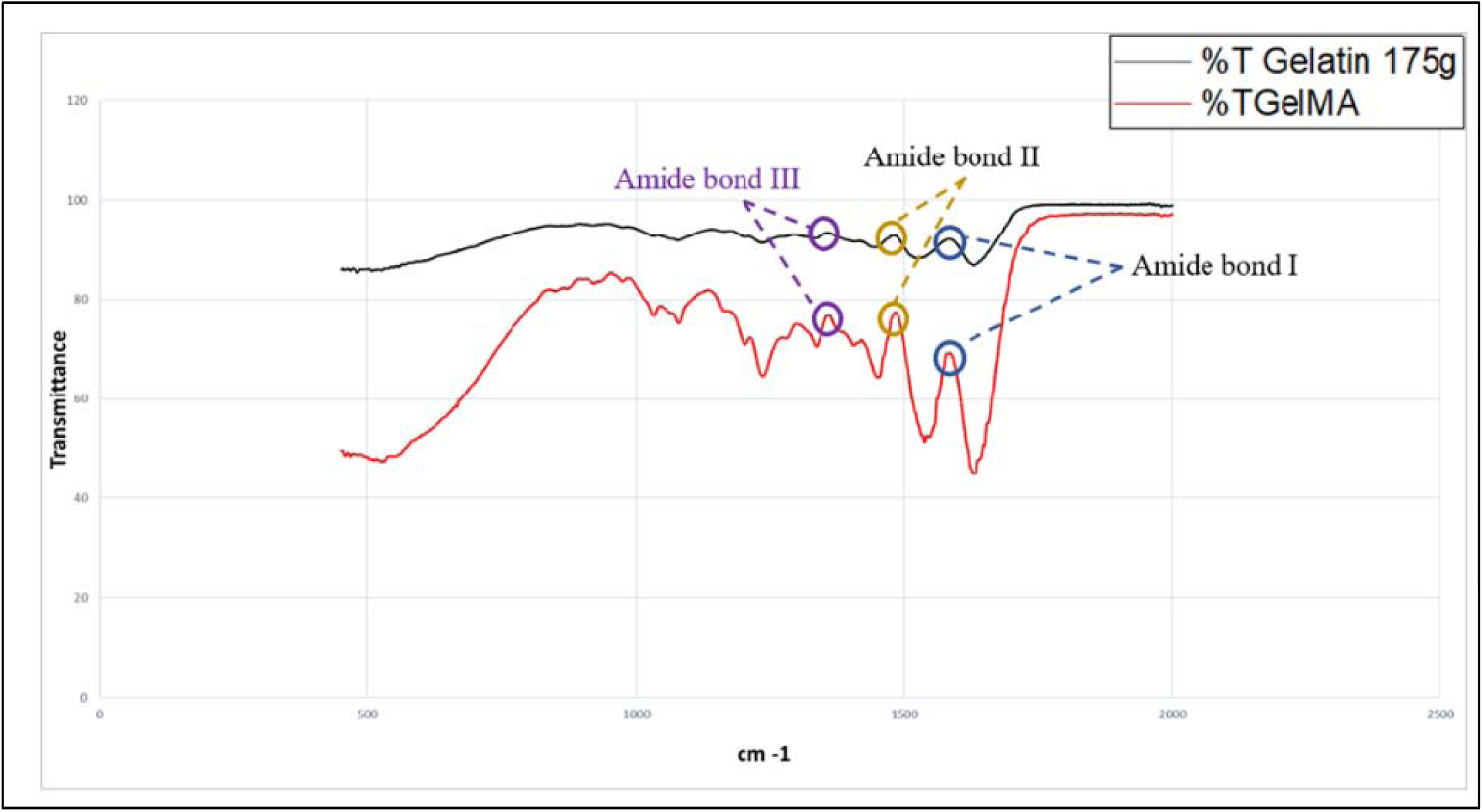
FTIR graph of gelatin and GelMA

#### 3.3.8 Decellularized ECM Solubilization

The SDS-PAGE was performed to determine the retention of the protein components in the solubilized dECM when compared with the extracted proteins from the native and dECM. **Figure 8B** demonstrates the SDS-PAGE results and the presence of bands in all the days of solubilization when compared to both native and dECM. The bands were found in varying sizes confirming the presence of all types of proteins mainly, type I collagen at around 120kDa and 250kDa. The total protein content was quantified by BCA assay and was found to be 1.28 ± 0.18 µg/mL for day 1, 1.39 ± 0.12 µg/mL for day 2, 1.12 ± 0.11 µg/mL for day 3, 1.19 ± 0.09 µg/mL for day 4, 1.4 ± 0.14 µg/mL for native and 1.3 ± 0.19 µg/mL for dECM (**Figure 8A**). The values are expressed as mean ± SD, with sample size of 04 on each day.

**Figure 8.**
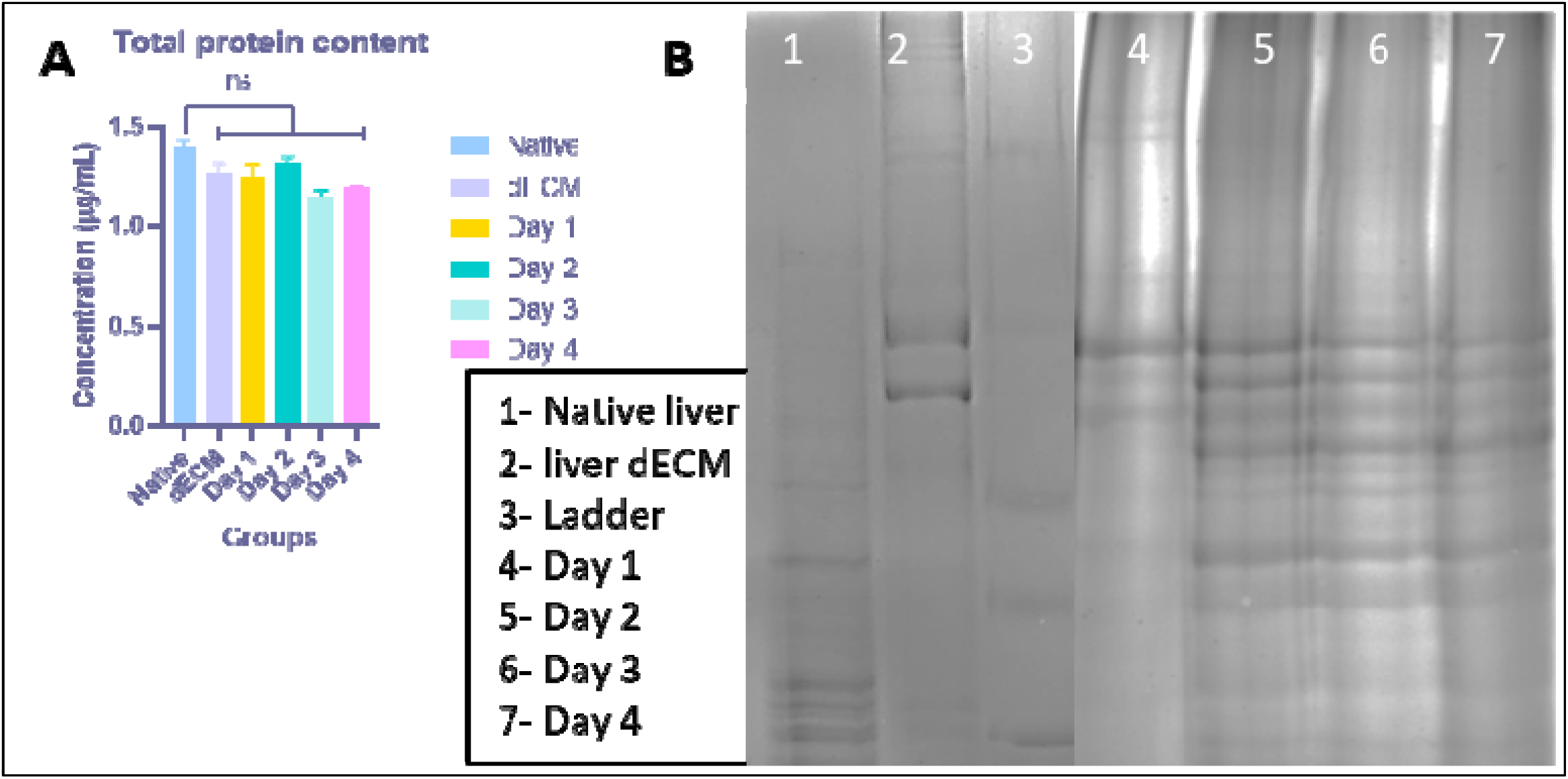
(A) Total protein concentration post solubilization on different days, (B) SDS-PAGE of solubilized dECM; (p<0.05) ns : non-significant

#### 3.3.9 Computer-Aided Design for Bioprinting

CAD models are digital designing through which the G-CODE to bioprint 3D models are generated (**Figure 9A**). The grid pattern shown corresponds to the infill or filament pathway, critical for determining the porosity and internal structure of the printed scaffold (**Figure 9B**). Actual photographic images of 3D-printed constructs (1x1x0.3 cm^3^) (**Figure 9C**). Close-up images show stacked, porous, lattice-like structures next to a ruler.

**Figure 9.**
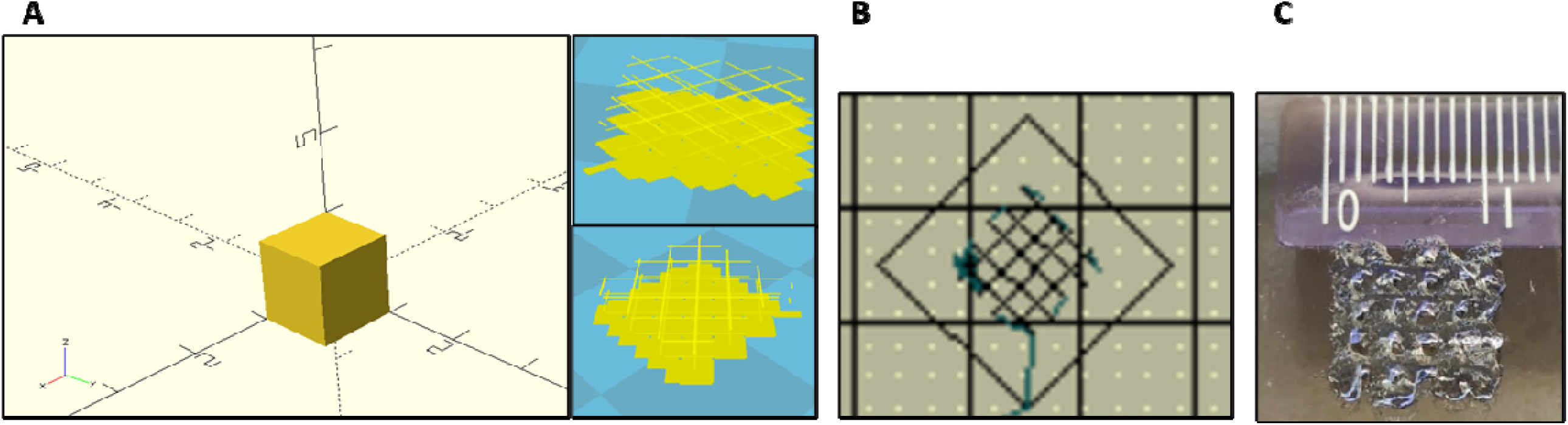
3D designing (A) CAD models designed, (B) G-CODE generated, and (C) 3D printed with GelMA

#### 3.3.10 Cytotoxicity Analysis of Solubilized dECM and GelMA on HepG2 Cells

The cytotoxicity of the ink was analysed by MTT assay on two different cell HepG2 and NIH3T3, to understand the effect of the ink on the viability of the cells. Both the cell lines showed good viability when compared to the control. It was found that the HepG2 cells had viability of 90% (**Figure 10A**) and NIH3T3 cells had viability of 93.8% (**Figure 10B**). The figure 10 represents the MTT graph of both the cells which proves that the ink is compatible with cells. Similarly, to confirm the cell viability of the cells exposed to the ink, scratch assay was performed on the NIH3T3 cells (**Figure 11**). After 30th hour the scratch in both the control and the test, the scratch was covered by the cells, confirming normal functioning of cells.

**Figure 10.**
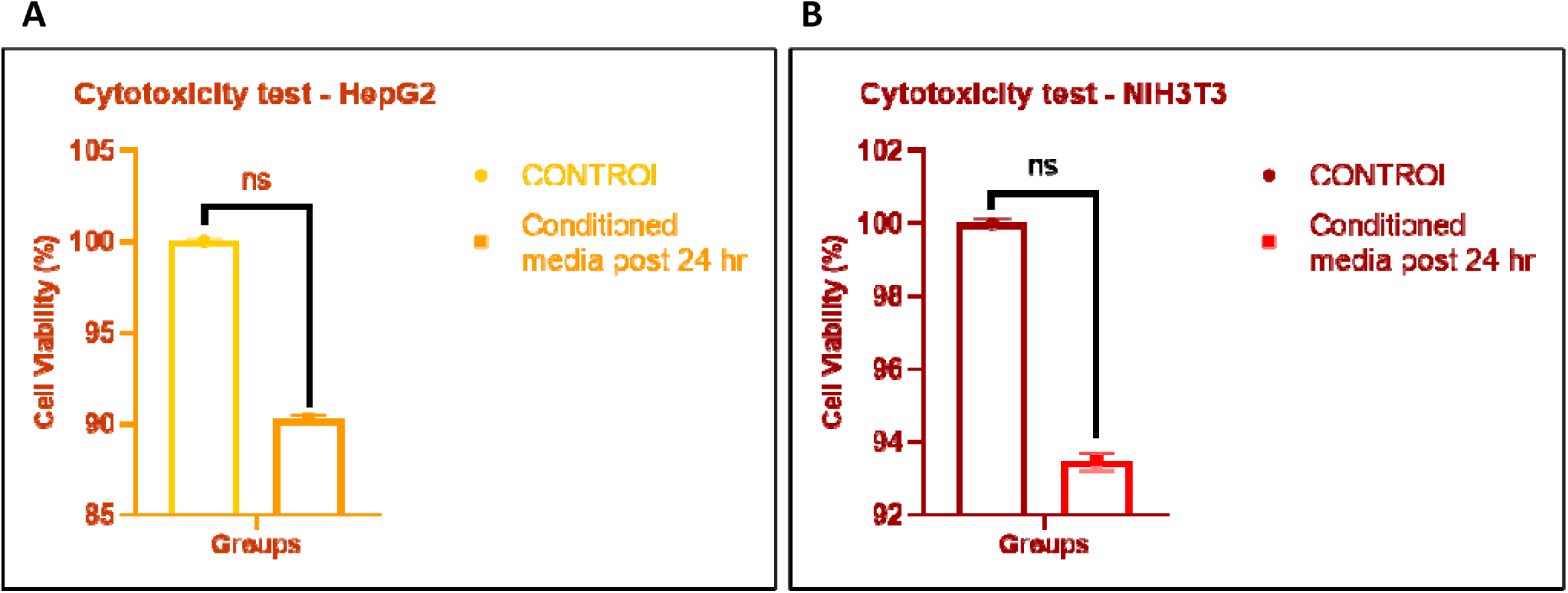
Cytotoxicity of ink on (A) HepG2 cells and (B) NIH3T3 cells; (p<0.05) ns : non-significant

**Figure 11.**
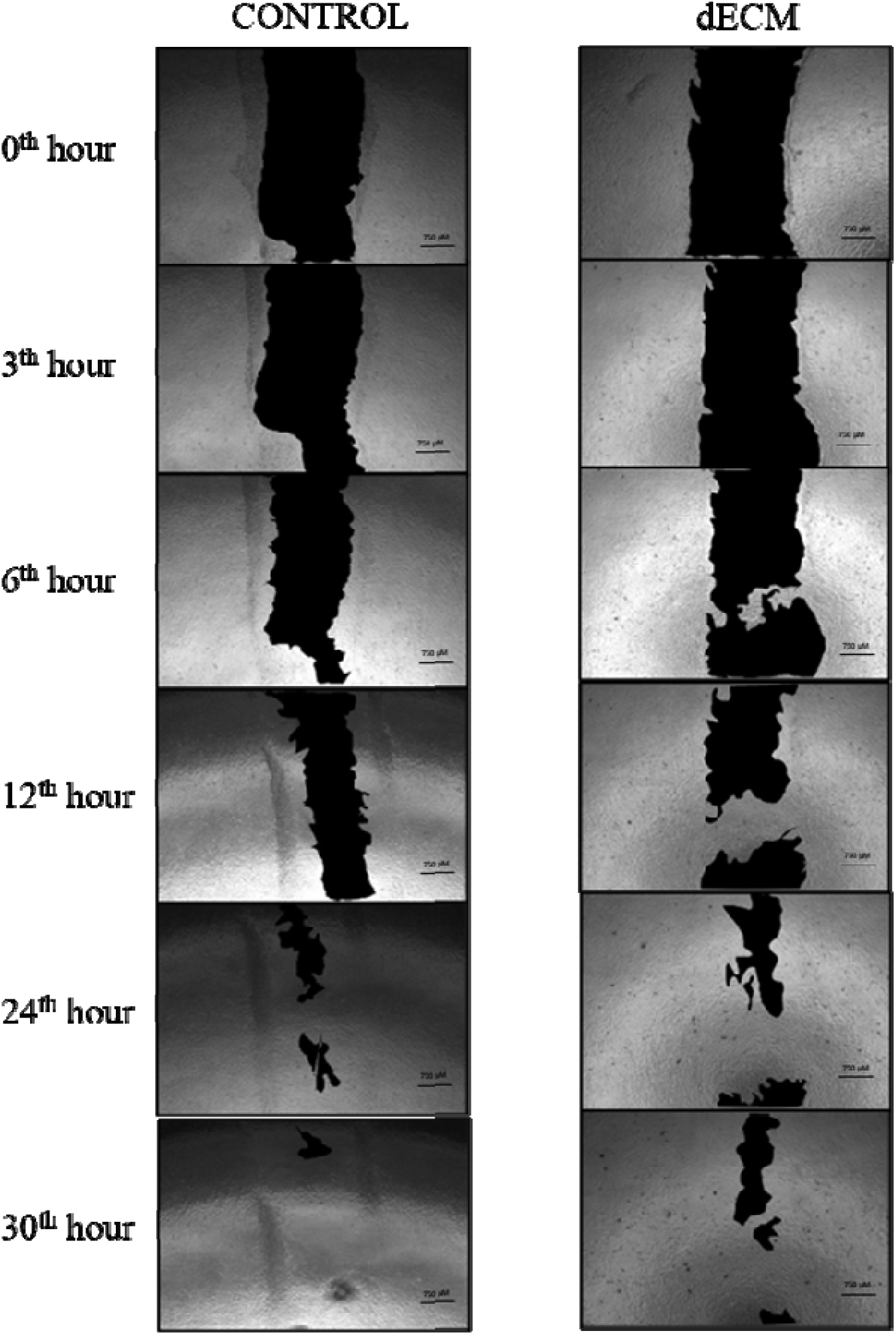
Scratch assay of ink (GelMA+dECM) on NIH3T3 cells

#### 3.3.11 Characterization of 3D Bioprinted Constructs

##### 3.3.11.1. Degradation Study

Degradation graph demonstrates 3D printed scaffold composition exposed to PBS and culture media (**Figure 12**) play critical roles in determining degradation kinetics, which are essential parameters for scaffold design in regenerative medicine and tissue engineering applications. Both groups show increased degradation over time, but scaffolds when subjected to culture media degrade faster than in PBS, at day 21, scaffolds in media reach a degradation rate of about ∼23%, and in PBS ∼16%. The enhanced degradation in media may be due to additional biological and enzymatic components (serum proteins) that are absent in PBS, facilitating faster breakdown of scaffold material.

**Figure 12.**
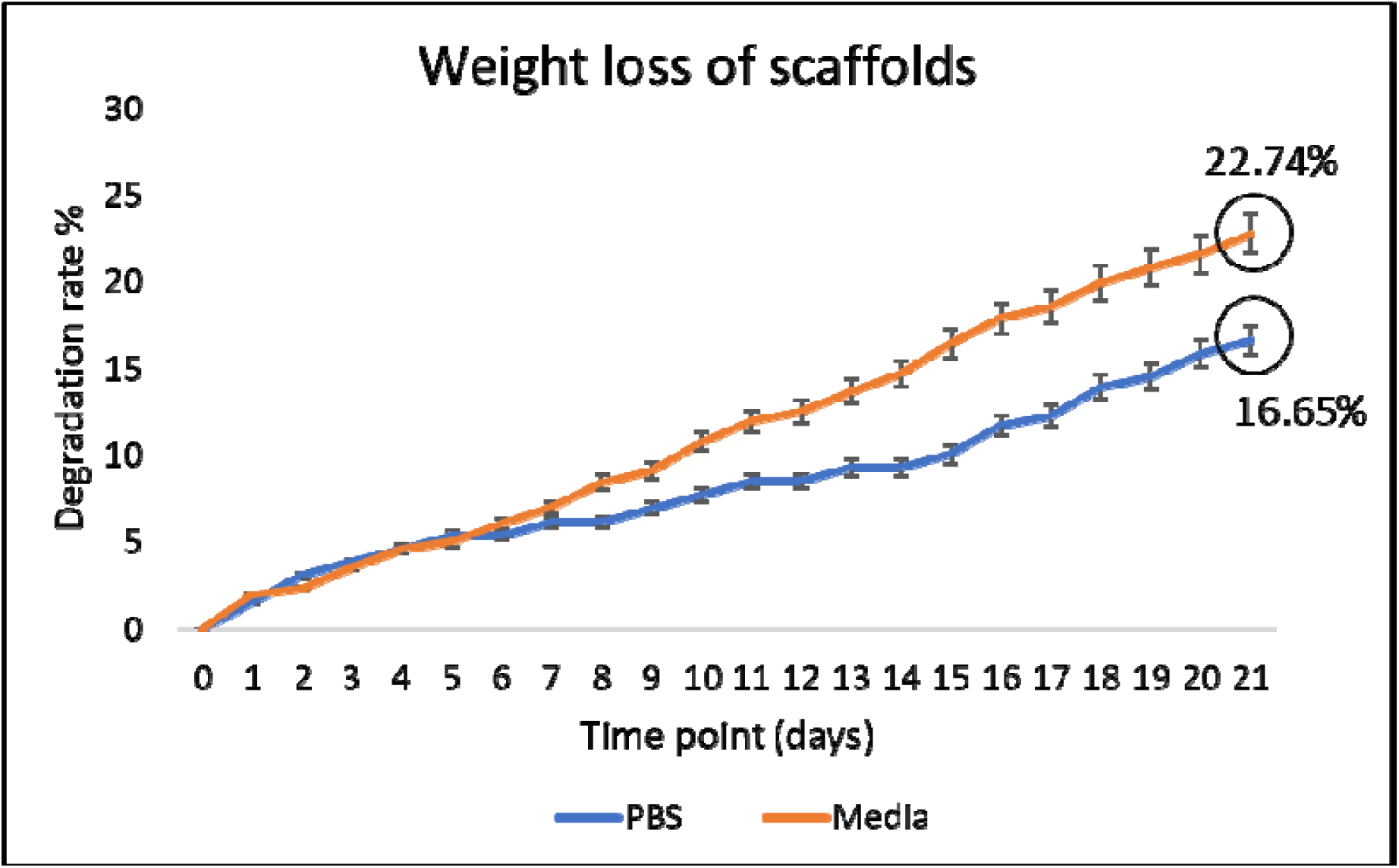
Degradation rate of scaffolds for 3 weeks in PBS and cell culture medium

##### 3.3.11.2. Induction of Fibrosis

Fibrotic conditions were induced in HepG2 cells cultured on bioprinted constructs by exposure to 10 mM methotrexate, optimized from previous studies, over a 72-hour period. Post-treatment, cell viability was analysed by MTT assay, and morphological evidence of fibrosis was evaluated using H&E staining. Cells were fixed in formaldehyde, sequentially dehydrated using graded ethanol washes, stained, and mounted with DPX prior to microscopic examination.

##### 3.3.11.3. Live/Dead Viability Assay

A Live/Dead viability assay was conducted on 3D bioprinted fibrotic scaffolds at Day 3. Scaffolds incubated with MTX exhibited an increased proportion of dead cells, as visualized by Z-stack merged fluorescence images. In contrast, untreated scaffolds demonstrated a predominance of viable cells relative to dead cells, whereas MTX-treated scaffolds exhibited the opposite trend, with a higher number of dead cells observed (**Figure 13**). The increased cell death in the fibrotic scaffolds following MTX treatment confirms the cytotoxic effect of the drug within the 3D bioprinted model.

**Figure 13.**
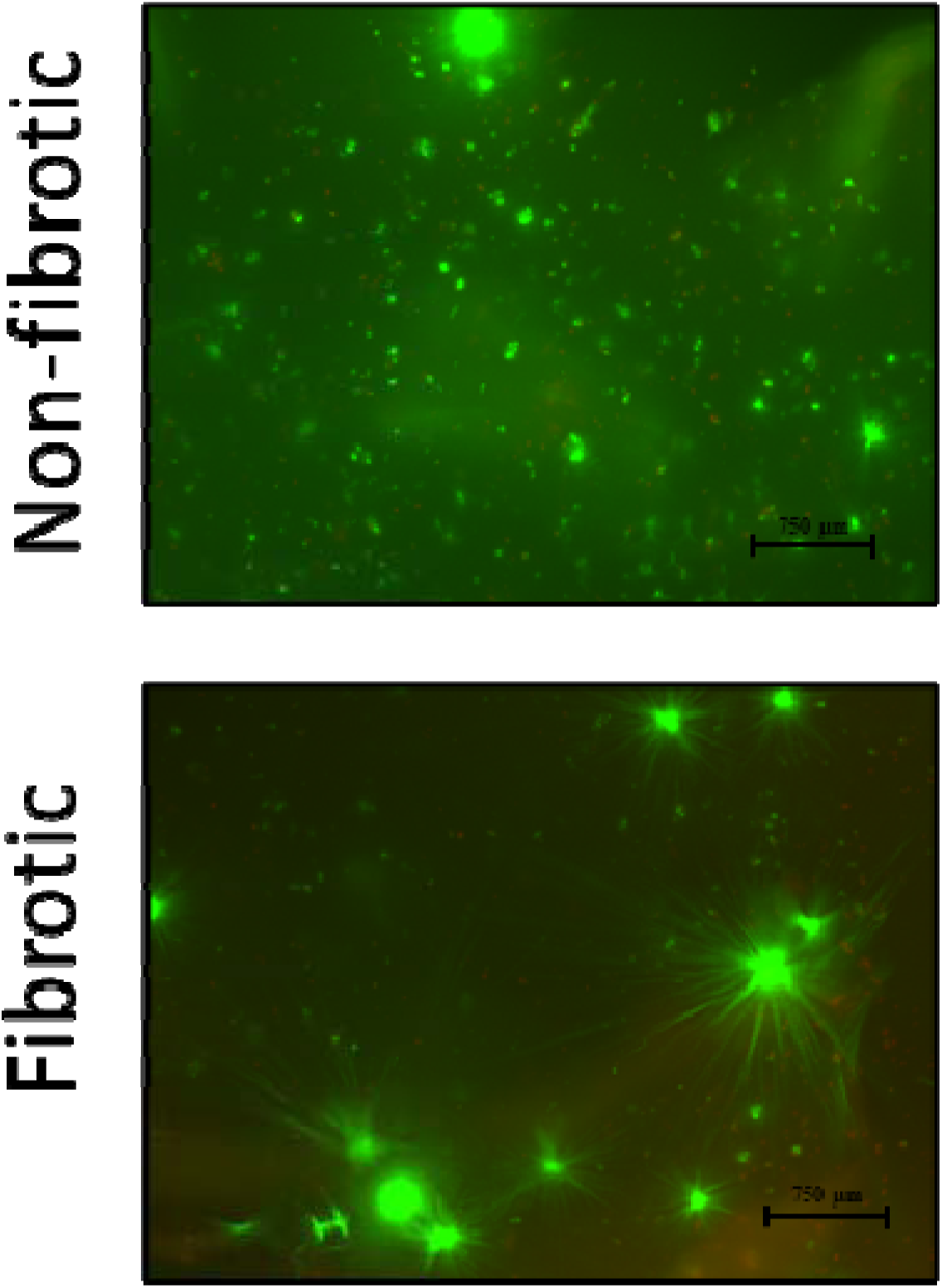
Live/ dead assay of 3D bioprinted non-fibrotic and fibrotic scaffolds (10X magnification)

##### 3.3.11.4. Histological Analysis of 3D Bioprinted Constructs

The H&E-stained images (**Figure 14B)** demonstrate the structural transition of bioprinted liver scaffolds upon fibrotic stimulation with MTX, confirming successful fibrosis induction *in vitro*. The switch from round to spindle-shaped cells indicates hepatic stellate cells (HSC) activation, a hallmark of liver fibrosis. Moreover, the increased eosin staining and altered morphology correlate with early fibrotic remodelling. The cells in non-fibrotic sections appear round-shaped, dispersed, and maintain a typical hepatic morphology with no visible architectural distortion or non-myofibroblast-like and Ishak score being 0, indicating no fibrosis (**Figure 14A**). The cells in fibrotic sections appear spindle-shaped morphology, contractile morphology, a hallmark of myofibroblast transdifferentiating and contractile morphology indicating activation of HSCs with no visible architectural distortion or non-myofibroblast-like and Ishak score being 3 along with dense eosinophilic regions, suggesting ECM accumulation, one of the classic features of fibrosis (**Figure 14B**). Ishak scoring assess the severity and progression of liver fibrosis in clinical and research settings. (**Figure 14C**) comprises of table describing the Ishak Staging System for liver fibrosis which is used in histopathology to grade the extent of fibrosis in liver biopsy samples. The table has two columns: Ishak score that lists stages from 0 to 6 and histological features describes the specific histological findings in the liver for each stage [38, 39].

**Figure 14.**
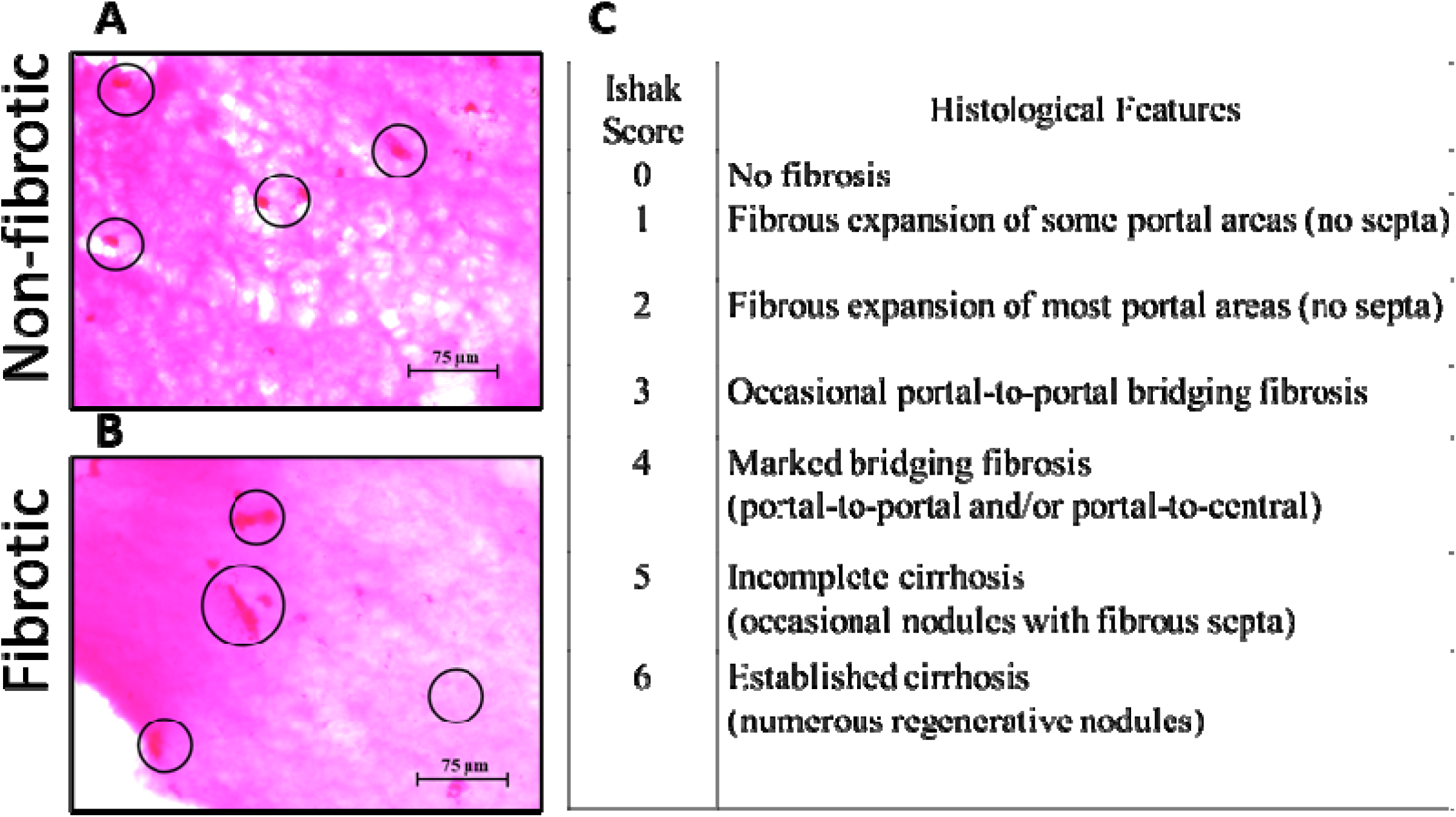
H&E staining of 3D bioprinted (A) non-fibrotic and (B) fibrotic scaffolds (40X magnification), (C) Ishak scoring [37]

##### 3.3.11.5. Functional Analysis of 3D Bioprinted Constructs

Functional evaluation of 3D bioprinted constructs under non-fibrotic (non-treated) and fibrotic (treated) conditions was performed by quantifying the release of liver-specific biochemical markers (**Figure 15**). Non-fibrotic constructs exhibited significantly higher concentrations of albumin, urea, and LDH compared to fibrotic constructs. Conversely, ALP) and ALT were significantly elevated in fibrotic constructs compared to controls. These findings suggest altered hepatic functionality consistent with induced fibrotic pathology, confirming the scaffold’s responsiveness to biochemical changes associated with liver fibrosis.

**Figure 15.**
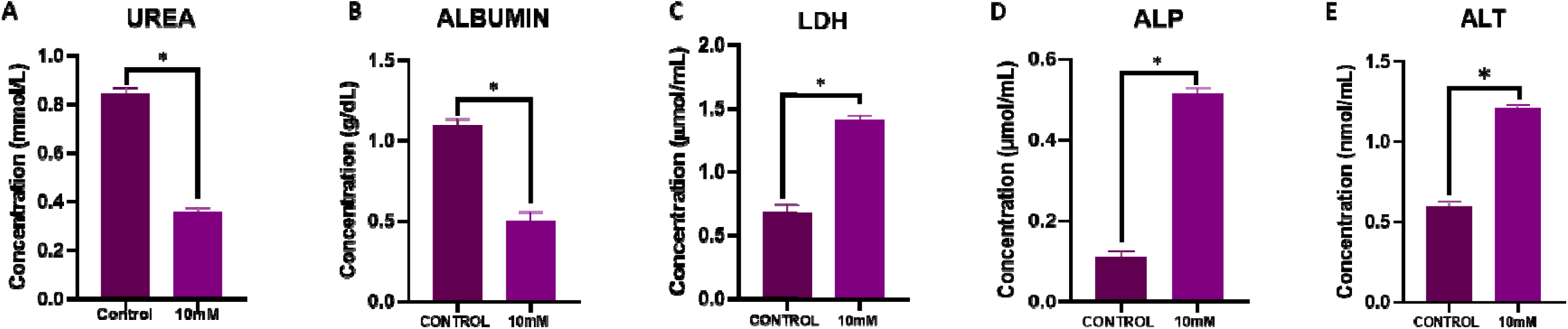
Functional tests on non-fibrotic and fibrotic scaffolds; * : statistically significant (p<0.05)

##### 3.3.11.6. Gene Expression Analysis

Quantitative real-time PCR was conducted to assess gene expression profiles in non-fibrotic and fibrotic 3D bioprinted constructs. Gene expression levels were normalized against the housekeeping gene GAPDH and presented as relative fold changes (**Figure 16**). Expression levels of hepatic markers (Cytochrome 3A4, Cytochrome 2E1, Cytochrome 2C9 and Albumin) and fibrosis-associated marker (Collagen I, Alpha-smooth muscle actin and Tissue inhibitor of matrix metalloproteinase 1) were significantly upregulated in fibrotic constructs compared to non-fibrotic controls. Elevated expression of these genes in the treated constructs is indicative of successful induction and manifestation of hepatic fibrosis at the molecular level, thus validating the utility of this bioprinted model for studying fibrotic mechanisms.

**Figure 16.**
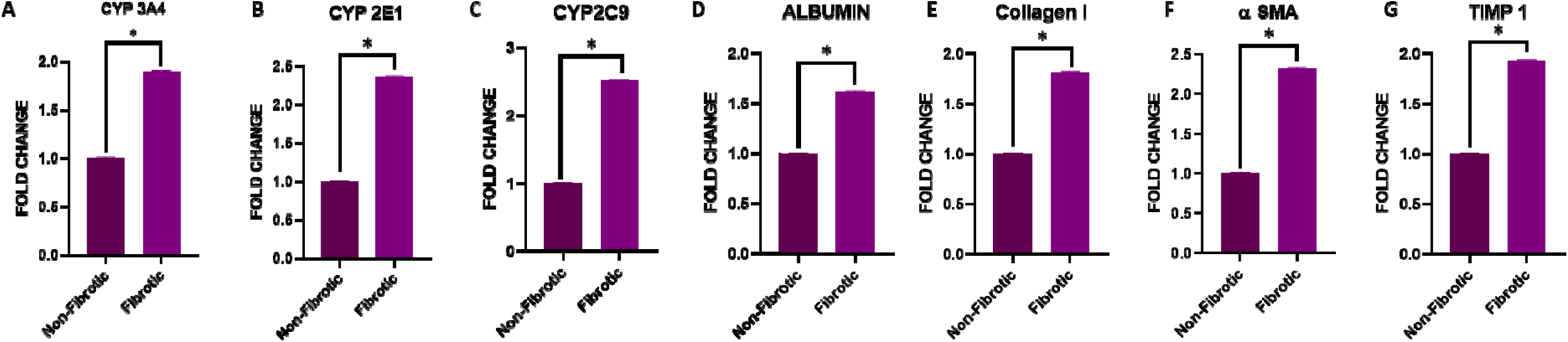
Gene expression on non-fibrotic and fibrotic scaffolds; * : statistically significant (p<0.05)

## 4.0 CONCLUSION

This study demonstrates a robust methodology for the decellularization of rat liver tissues, GelMA synthesis, and development of a biocompatible GelMA-dECM bioink for 3D bioprinting in hepatic tissue engineering. The decellularization process efficiently preserved essential ECM proteins, GAGs, and scaffold integrity, while GelMA provided the structural stability necessary for scaffold formation and sustained cellular viability and proliferation. Functional and molecular validations confirmed that the 3D bioprinted constructs effectively mimic non fibrotic and methotrexate-induced fibrotic liver conditions. This innovative model holds substantial translational promise, specifically as a powerful platform for high-throughput drug screening, toxicity assessment, and precision modeling of liver fibrosis, thereby potentially accelerating therapeutic advancements in hepatic diseases.

## ACKNOWLEDGEMENTS

The authors wish to acknowledge the Indian Council for Medical Research for funding the research. The authors would like to thank the Manipal Center for Biotherapeutics Research (MCBR), MAHE, and Manipal for the provision of research instruments and infrastructure support.

## 6. AUTHOR’S CONTRIBUTION

Kirthanashri S. V.: Conceptualization, Investigation, Data Curation, Writing – original draft, editing, reviewing and validation; Mrunmayi Gadre: Investigation, Data Curation, Visualization, Writing – original draft

## **7.** AVAILABILITY OF DATA AND MATERIALS

The datasets generated and/or analysed during the current study are available online as the Indian patent, repository (Indian Patent No. 202341045531)

